# Can Zika account for the missing babies?

**DOI:** 10.1101/126557

**Authors:** Flávio Codeço Coelho, Margaret Armstrong, Valeria Saraceni, Cristina Lemos

## Abstract

The Zika virus (ZIKV) spread rapidly in Brazil in 2015 and 2016. Rio de Janeiro was among the Brazilian cities which were hit the hardest, with more that a hundred thousand confirmed cases up to the end of 2016. Given the severity of the neurological damage caused by ZIKV on fetuses, we wondered whether it would also cause an increase in the number of miscarriages, especially very early ones. As early miscarriages are unlikely to be recorded as a health event, this effect – if it occurred – would only show up as a reduction in the number of live births. In this paper we show that there was a 15% drop in live births between September and December 2016 compared to the previous year, and that this sharp drop from epidemiological week 33 onward is strongly correlated with the number of recorded cases of Zika about 40 weeks earlier. We postulate that ZIKV is directly responsible for this drop in the birth rate. Further work is required to ascertain whether other factors such the fear of having a microcephaly baby or the economic crisis are having a significant effect.

## 1. Introduction

Propelled by a combination of vectorial and sexual transmission, the Zika virus (ZIKV) spread rapidly in Brazil in 2015 and 2016, with the epidemic reaching neighboring countries very quickly[4]. Rio de Janeiro was among the Brazilian cities which were hit the hardest. With more than 6 million inhabitants, it is an excellent place to observe effects which would be hard to detect in smaller populations. It was based on data from Rio that the increased risk of Zika in females of reproductive age, was first described [2].

The data on the Zika epidemic in Rio still keeps revealing important new facts about this emerging infection. In a recent publication, de Oliveira et al. [3], looking at the cases of Zika, Guillain-Barreé Syndrome (GBS) and microcephaly commented that the data collected cannot explain why there are many fewer cases of microcephaly than expected in 2016, whereas the number of cases GBS was more or less as expected. They propose three possible reasons for this: 1) the increase in the cases of GBS was due to another virus such as CHIKV rather than ZIKV, and so cases of fever were wrongly attributed to ZIKV; 2) ZIKV is a necessary but not sufficient cause for microcephaly; 3) fear of adverse consequences led to fewer conceptions.

In this paper we propose a fourth possibility: ZIKV causes miscarriages early in pregnancy – even before the mother realizes that she is pregnant. This hypothesis is supported by recent reports about early miscarriages related to ZIKV infection [7, 1]. A report by PAHO points out that out of 582 fetal deaths, 200 were confirmed as central nervous system malformations[5].

Using data on live births from the city of Rio de Janeiro up to the end of 2016 we show that 1) there were 5154 fewer births in the second half of 2016 (7484 for the whole year) compared to previous years (i.e. a 14.85% drop in the number of births) and 2) this drop is correlated with the number of ZIKV cases.

As early miscarriages are unlikely to be recorded as a health event, this effect – if it occurred – would only show up as a reduction in the number of live births.

In Brazil, there is a a fairly complete and reliable record of live births. As births in Rio de Janeiro show seasonality, we compare the number of live births year on year for a given epidemiological week. Figure 1 shows the number of live births per week in 2016 in red compared to the average number per week from 2012 to 2015 in blue with the 95% confidence shaded around it. From week 33 onward (i.e. about 40 weeks after the start of the ZIKV epidemic) the red line is consistently below the confidence interval.

**Figure 1:**
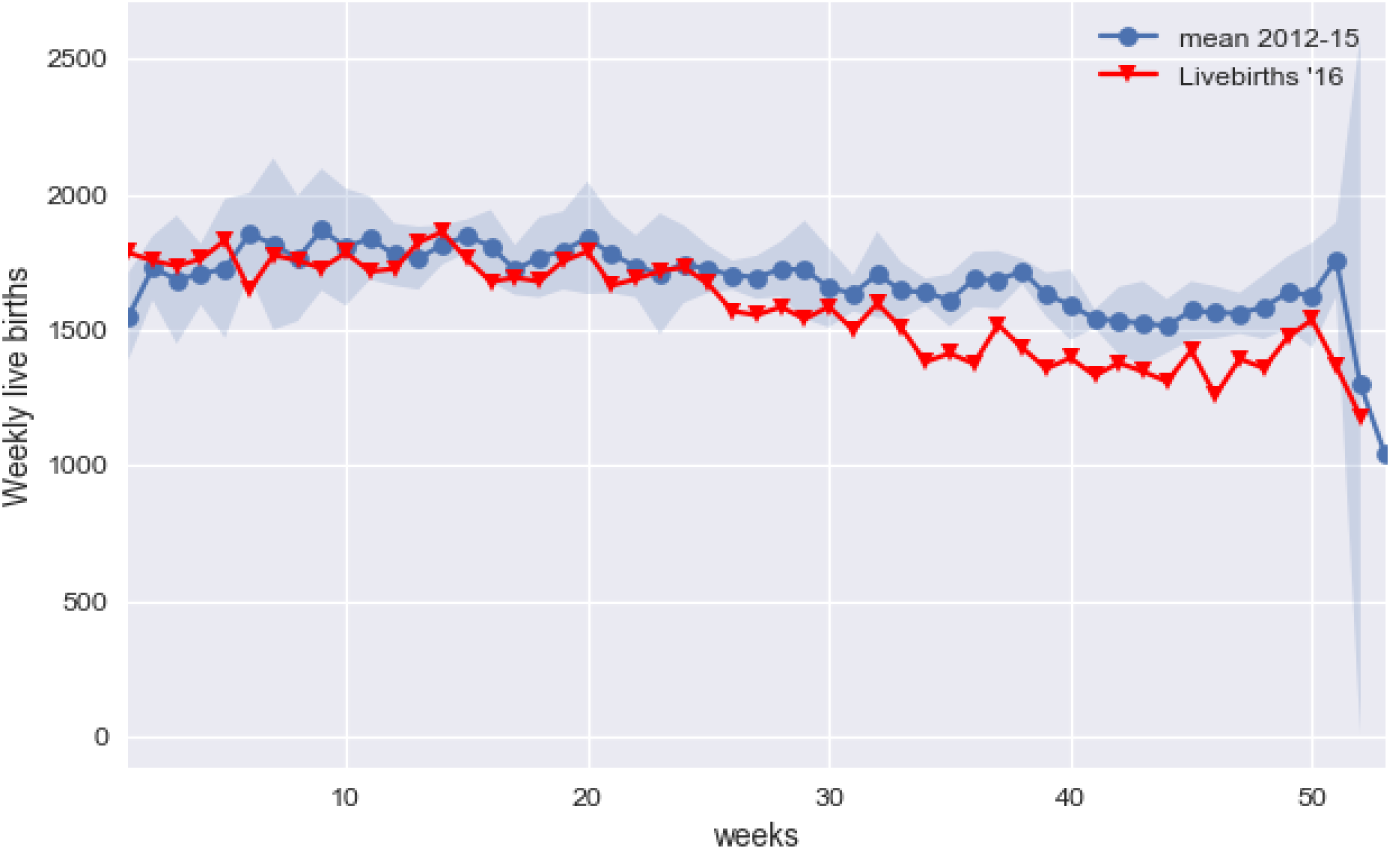
Average births per week for year 2012-15 (in blue) and weekly births in 2016.

Moreover the drop in births shows a significant positive correlation with the incidence of Zika between 37 and 42 weeks earlier(figure 2) which corresponds to the typical human pregnancy duration. To further strengthen the hypothesis of causal association between the Zika incidence and the drop in births we also included the incidence of Chikungunya in the model as well.

**Figure 2:**
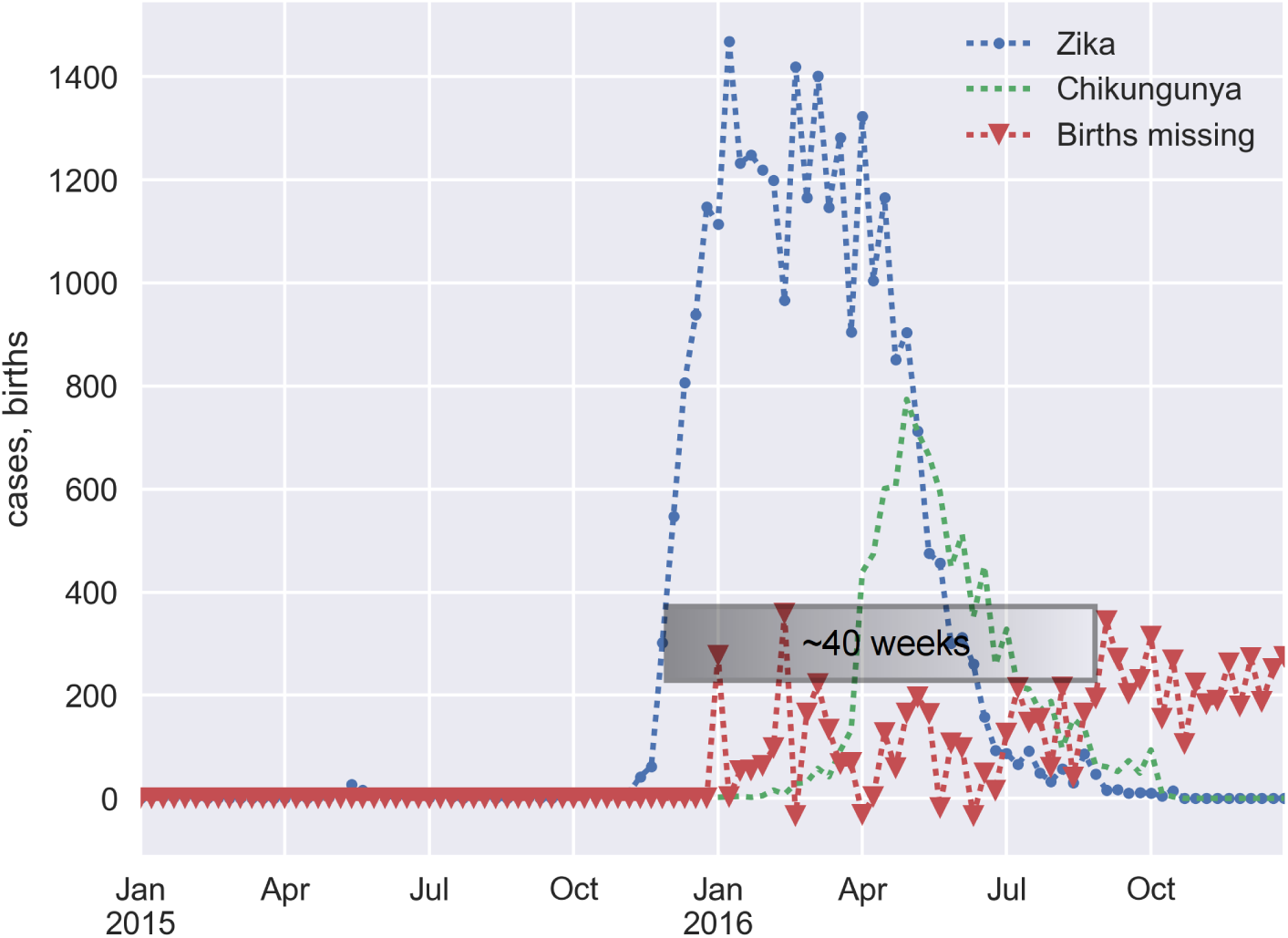
The lag between the onset of the zika outbreak and the increase in the loss of births is roughly 40 weeks.

## 2. Methods

Data about live births from the city of Rio de Janeiro was obtained from SINASC, the national system for live births registration. The birth rate in Rio has been stable for a number of years.

The reported cases of Zika were obtained from SINAN, the national registry of diseases of mandatory reporting. Reporting of Zika became mandatory in late 2015. We only had access to SINAN and SINASC data from the city of Rio de Janeiro city.

As births show a natural annual seasonality, we set out to measure the loss in births on a week-by-week basis. Let 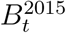 be the number of birth live births on week *t* of 2015. Let 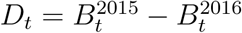 for *t* ranging from 1 to 52, the number of weeks in one year. Let *Z_t_* and *C_t_* be the number of female Zika and Chikungunya cases notified on week *t*, respectively. Assuming *D_t_* follows a Gaussian distribution, we proposed the following generalized linear model:

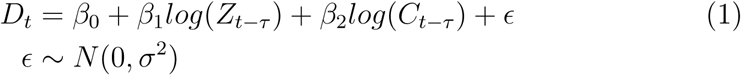
where *τ* has units of weeks and takes values from the interval (33, 45). This model tests the hypothesis that the incidence of Zika *τ* weeks earlier is positively correlated with the deficit of births in week *t*, defined above and denoted by *D_t_*.

## 3. Results

Taking the average number of births from 2012-15 as the expected number for 2016, there were 5154 missing babies, from week 33 on. This figure represents a 14.85% drop in expected number of births for this period. The number of missing babies for the entire year was 7484, so the bulk of the deficit in births really occurred after week 33.

The results in table 1, show that the weekly loss of births, is statistically associated with the incidence of Zika in the past. The association is stronger in the time window of 38 to 42 weeks (p < 0.01). Weaker but still significant (*p* < 0.05) association can be seen in the range of 35-37 and at 43 and 45 weeks after the Zika epidemic.

The incidence of Chikungunya does not seem to correlate at all with the drop in births for the same lag range.

**Table 1:** Results of fitting the model to the data for τ ranging from 33 to 45. Figures in bold are significant at *p* < 0.01.

| $\tau$ (weeks) | $\beta_1$ | $\beta_2$ | **p-values:** $\log(Z_{t-\tau}), \log(C_{t-\tau})$ |
| --- | --- | --- | --- |
| 33 | 15.56 | 6.83 | 0.11, 0.82 |
| 34 | 10.28 | 30.29 | 0.27, 0.30 |
| 35 | 21.93 | -16.45 | 0.02, 0.57 |
| 36 | 20.78 | -2.88 | 0.02, 0.91 |
| 37 | 17.36 | 8.54 | 0.05, 0.76 |
| **38** | **24.23** | -15.7 | **0.007**, 0.57 |
| **39** | **24.24** | -5.55 | **0.005**, 0.83 |
| **40** | **25.27** | -10.29 | **0.001**, 0.67 |
| 41 | 18.58 | 7.13 | 0.022, 0.77 |
| **42** | **29.24** | -34.96 | **0.000**, 0.162 |
| 43 | 17.99 | 7.74 | 0.03, 0.76 |
| 44 | 15.38 | 13.16 | 0.07, 0.62 |
| 45 | 23.94 | -47.85 | 0.025, 0.14 |

## 4. Discusion

Although the analysis presented does not completely rule out other influences on Rio de Janeiro’s birth rate, it demonstrates a strong correlation between the drop in the birth rate and the number of Zika cases. The actual range of lags of the influence and the lack of an association with Chikungunya, which happened at approximately the same period, points strongly to Zika being a causal factor in this loss of babies.

The implications of a ZIKV infection during pregnancy for the health of fetuses, is quite well established [6], but the implication of Zika on early miscarriages [7] could benefit from larger scale studies. Our results point to a potential huge loss of births in the rest of the country, but unfortunately the data for the whole country was not available to us.

We hope the evidence provided by this study will lead to more specific investigations to determine if miscarriages after a Zika epidemic can be traced back to infections during pregnancy. The drop in the birth rate, even if only due to family planning, would still be a serious consequence of a Zika epidemic, because of its economic implications.

## References

[1] Brasil, P., J. P. Pereira, C. Raja Gabaglia, L. Damasceno, M. Wakimoto, R. M. Ribeiro Nogueira, P. Carvalho de Sequeira, A. Machado Siqueira, L. M. Abreu de Carvalho, D. Cotrim da Cunha, and et al. 2016. Zika virus infection in pregnant women in rio de janeiropreliminary report. New England Journal of Medicine.

[2] Coelho, F. C., B. Durovni, V. Saraceni, C. Lemos, C. T. Codeo, S. Camargo, L. M. Carvalho, L. Bastos, D. Arduini, D. Villela, and et al. 2016. Sexual transmission causes a marked increase in the incidence of zika in women in rio de janeiro, brazil. bioRxiv, P. 055459.

[3] de Oliveira, W. K., E. H. Carmo, C. M. Henriques, G. Coelho, E. Vazquez, J. Cortez-Escalante, J. Molina, S. Aldighieri, M. A. Espinal, and C. Dye 2017. Zika virus infection and associated neurologic disorders in brazil. New England Journal of Medicine, 0(0).

[4] Fauci, A. S. and D. M. Morens 2016. Zika virus in the americasyet another arbovirus threat. New England Journal of Medicine.

[5] Organization, P. A. H. 2017. Zika - epidemiological report brazil. march 2017. washington. Technical report, PAHO/WHO.

[6] Teixeira, M. G., M. da Conceio N Costa, W. K. de Oliveira, M. L. Nunes, and L. C. Rodrigues 2016. The epidemic of zika virus-related microcephaly in brazil: Detection, control, etiology, and future scenarios. American Journal of Public Health, 106(4): 601605.

[7] van der Eijk, A. A., P. J. van Genderen, R. M. Verdijk, C. B. Reusken, R. Mgling, J. J. van Kampen, W. Widagdo, G. I. Aron, C. H. GeurtsvanKessel, S. D. Pas, and et al. 2016. Miscarriage associated with zika virus infection. New England Journal of Medicine, 375(10): 10021004.

